# Vomocytosis of *Cryptococcus neoformans* cells from murine, bone marrow-derived dendritic cells

**DOI:** 10.1101/2022.03.28.486165

**Authors:** Noah Pacifici, Melissa Cruz-Acuña, Agustina Diener, Neeraj Senthil, Hyunsoo Han, Jamal S. Lewis

## Abstract

*Cryptococcus neoformans* (CN) cells survive within the acidic phagolysosome of macrophages for extended times, then escape without impacting the viability of the host cell via a phenomenon that has been coined ‘vomocytosis’. Through this mechanism, CN disseminate throughout the body, sometimes resulting in a potentially fatal condition - Cryptococcal Meningitis (CM). Justifiably, vomocytosis studies have focused primarily on macrophages, as alveolar macrophages within the lung act as first responders that ultimately expel this fungal pathogen. Herein, we hypothesize that dendritic cells (DCs), an innate immune cell with attributes that include phagocytosis and antigen presentation, can also act as ‘vomocytes’. Presciently, this report shows that vomocytosis of CN indeed occurs from DCs. Primarily through time-lapse microscopy imaging, we show that rates of vomocytosis events from DCs are comparable to those seen from macrophages and further, are independent of the presence of the CN capsule and infection ratios. Moreover, phagosome-altering drugs such as chloroquine and bafilomycin A, as well as the actin-modifying drug, cytochalasin B inhibit this phenomenon from DCs. Although DC immunophenotype does not affect the total number of vomocytic events, we observed differences in the numbers of CN per phagosome and expulsion times. Interestingly, these observations were similar in primary, murine macrophages. Understanding the vomocytic behavior of different phagocytes and their phenotypic subtypes is needed to help elucidate the full picture of the dynamic interplay between CN and the immune system. Critically, deeper insight into vomocytosis could reveal novel approaches to treat CM, as well as other immune-related conditions.

## Introduction

Pathogens have evolved over time to develop means for persistence, dissemination, and infection within their mammalian hosts. The fungal species *Cryptococcus neoformans* (CN) is an opportunistic pathogen that causes an infectious disease called ‘Cryptococcosis’ that predominantly affects immunocompromised patients—primarily those afflicted with HIV/AIDS^1–3^. This infection can deteriorate into a condition called Cryptococcal Meningitis (CM), where CN establishes an infection in the central nervous system (CNS). Globally, there are estimated to be 223,100 CM cases and 181,000 deaths annually^1^.

*Cryptococcus neoformans* spores typically enter the host through the lung via inhalation. Subsequently, the fungal pathogen disseminates from the lung into other tissues, including the CNS^2,4^. The three proposed mechanisms that facilitate CN entry into the CNS are paracytosis, transcytosis, and hitchhiking. In the latter, CN are suggested to cross the blood brain barrier (BBB) by ‘hitchhiking’ within host phagocytes^5,6^. This is known as the ‘Trojan Horse’ hypothesis. *Kechichian et al*. showed that depletion of alveolar macrophages in mice significantly reduces cryptococcal dissemination to the CNS^7^. Later, *Charlier et al*. showed that infecting naïve hosts with CN-infected monocytes significantly increases CN accumulation in the brain, compared to infecting with CNs directly^8^.

Further, Cryptococcal cells have been shown to escape from macrophages (MΦs) by inducing expulsion whilst leaving the phagocyte unharmed through a phenomenon called ‘vomocytosis’ (non-lytic exocytosis)^3,9–11^.

Like most cell types, macrophages regularly perform exocytosis to recycle membrane components and excrete various factors^12–14^. However, vomocytosis is a unique form of exocytosis whereby these immune cells slowly (over 5 to 20 hours) expel large pathogenic particulates that they are otherwise programmed to be retained and digested. Studies have identified intracellular, physicochemical, and immunological cues in CN-infected MΦs that are linked to this mechanism. As MΦs perform vomocytosis, actin rapidly and transiently polymerizes in a cage-like structure around the phagosome, which then fuses with the plasma membrane^15^. These cages may be a post-phagosome permeabilization attempt by the MΦ to inhibit CNs’ escape, as inhibiting actin polymerization has been shown to increase vomocytosis occurrence. In addition, CN disrupts phagolysosomal maturation, as characterized by the rapid removal of the early phagosome markers Rab5 and Rab11^16^. Some reports have suggested that there is alkalinization of, and abnormal calcium ion levels in phagosomal compartments that contain the live pathogen^16–20^. Furthermore, the addition of weak bases to CN-infected macrophages has been shown to modulate vomocytosis occurrence from macrophages^17,20^. Moreover, *Gilbert et al*. showed that pharmacological inhibition of ERK5 increases vomocytosis occurrence^21^.

Further, the immune state of infected MΦs has been suggested to also influence this phenomenon. A recent study by *Seoane et al*. discovered that viral exposure to either measles or human immunodeficiency virus (HIV) were both shown to significantly boost expulsion rates of CN cells from MΦs^22^. Moreover, other factors known to elicit an antiviral response—the TLR3 agonist poly(I·C) and type I interferons, IFN-α and IFN-β—all similarly increase MΦ vomocytosis events. Another study investigated how different T cell effector-induced phenotypes can impact the expulsion rates of infected J774 MΦs^23^. Prior to infection, these cells were treated with cytokines inducing T effector cell-induced phenotypes Th1 (IFN-γ and TNF-α), Th2 (IL-4 and IL-13), or Th17 (IL-17). The Th1 and Th17 subtypes of J774 cells showed diminished intracellular CN proliferation and increased vomocytosis rates, whilst the Th2 group displayed increased intracellular CN proliferation and reduced vomocytosis occurrence.

Taken altogether, studies have given some clarity on the influence of intracellular, physicochemical, and immune states on vomocytosis. However, they focus entirely on MΦs, which is only a single cell type in an army of immune cells. Moreover, *Yang et al*. recently discovered the occurrence of this phenomenon in neutrophils^24^.

Like macrophages and neutrophils, dendritic cells (DCs) have the unique ability to phagocytose particulates, including pathogens. More importantly, DCs link the innate and adaptive arms of the immune system. These innate immune cells are key for maintaining a coherent dynamic on both, the host defense against pathogens and protection of “self” antigens of host cells and tissues^25–27^. Dendritic cells detect invading pathogens due to their constituent sensors (e.g. Toll-like receptors [TLRs])^28,29^. They communicate the presence of pathogens to the adaptive immune system, thereby initiating long lasting, antigen-specific responses. Migration of DCs to T cell-rich regions is critical here and is mainly regulated by the chemokine receptor CCR7^30–32^ and CCL21^33–35^. Following DC migration to secondary lymphoid organs, lymphocytes are subsequently activated and induced to proliferate and become potent effector cells (e.g. helper T cells)^25^. Interestingly, other innate immune cell types can traffick via lymphatics^32,36^ and perform antigen presentation^37–39^. While these cell types have some promising capability in lymphatic migration and antigen presentation, their abilities are limited in comparison to DCs, which are recognized as the primary antigen presenting cell type whose dominant function is to traffick to the LNs to present foreign material to LN-resident T cells.

During cryptococcal infections, DCs are known interact with CN cells and have been demonstrated to phagocytose CN cells following opsonization with complement or antibody^40^. *Hole et al*. demonstrated the ability of DC lysosomal extract to cause morphological changes in CN and kill the pathogen *in vitro* via oxidative and non-oxidative mechanisms^41^. *Artavanis-Tsakonas et al*. showed that CN-containing DC phagosomes have an impaired CD63 recruitment, indicative of a distinct phagosomal compartment composition that may affect the outcome of antigen processing and presentation^42^. However, to the best of our knowledge, the ability of DCs to expel CN from their phagosome via vomocytosis has not been investigated thus far.

We hypothesized that DCs could perform vomocytosis, especially given that they share much of the same vacuolar machinery as their phagocytic relatives, MΦs and neutrophils. Herein, we endeavored to document vomocytosis from DCs— a key player in bridging the innate and adaptive immune response. Further, we investigated the effect of CN infection ratio, presence of CN capsule, drug manipulation of the phagosome conditions and actin polymerization on vomocytosis from DCs. The overall effect of the immune state on vomocytosis from both DCs and MΦs was also assessed. Finally, we characterized vomocytosis based on multiple outcomes— rate of vomocytosis occurrence, timing of expulsion, and number of internalized CN prior to expulsion. We believe that further investigation of CN’s complex interactions with different phagocytic cell types will act as a step towards elucidating the complex story of cryptococcal infections and underlying mechanisms of vomocytosis.

## Materials and Methods

### Bone marrow-derived Dendritic Cell and Macrophage (MΦ) Culture

Primary DCs and MΦs were obtained from the bone marrow of C57BL/6 mice as described in previous studies^43^. Growth media consisted of DMEM/F-12 1:1 with L-glutamine (Cellgro, Herndon, VA), 10% fetal bovine serum, 1% sodium pyruvate (Lonza, Walkersville, MD), 1% nonessential amino acids (Lonza, Walkersville, MD), 1% penicillin/streptomycin (Cytiva, Marlborough, MA) and 20 ng/mL GM-CSF (R&D Systems, Minneapolis, MN) (DC media) or L929 conditioned media (MΦ media) and incubated at 37°C and 5% CO_2_. For conciseness, media containing all necessary growth factors and added reagents for the necessary cell type will be denoted as ‘complete media’. Unless otherwise noted we used DC and MΦ on day 10 of their respective cultures

### Cell Phenotype Validation via Flow Cytometry

On day 6 of DC or MΦ culture, cells were characterized by measuring the presence of phenotype-specific surface markers, with antibodies against F4/80 [APC, BM8 Clone] (eBioscience, San Diego, CA) and CD11c [PE-Cy7, HL3 Clone] (BD Pharmingen, San Diego, CA) via an Attune Nxt Flow Cytometer (Life Technologies, Carlsbad, CA).

### Dendritic Cell and MΦ Polarization Validation

Dendritic cells and MΦs were seeded on 12 well plates (6 million cells/ plate [DCs] and 3 million cells/ plate [MΦs]) in complete media containing polarizing agents. The difference in seeding density was due to spreading ability, with MΦs being much more elongated than DCs and therefore taking up more surface area per cell. For inflammatory activation, cells were treated with LPS (100 ng/ml) from *Escherichia coli* O111:B4 (Sigma, St. Louis, MO). For anti-inflammatory polarization DCs were treated with 1uM of dexamethasone (DEX; Alfa Aesar, Tweksbury, MA) and MΦs with IL-4 (20ng/ml) and IL-13 (20ng/ml; denoted ‘IL4/13’ for brevity; R&D Systems) in MΦ media. After a 48-hour incubation period, the expressions of cell surface markers were determined via flow cytometry using antibodies against F4/80, CD38 [PerCP-eFluor 710, 90 Clone] (eBioscience), Arginase 1 [PE, A1exF5 Clone] (eBioscience), and iNOS [PE-Cy7, CXNFT Clone] (eBioscience) for MΦs and antibodies against CD11c, MHCII [Alexa-Fluor 488, M5/114.15.2 Clone] (BD Pharmingen), CD80 [APC, 16-10A1 Clone] (BioLegend, San Diego, CA), and CD86 [PE, GL1 Clone] (BD Pharmingen) for DCs. Additionally, an LPS activation resistance test was performed by adding LPS (100 ng/ ml) to DCs and MΦs previously treated with tolerogenic polarization agents for an additional 48 hours. Subsequently, the expressions of the same cell surface markers were quantified using flow cytometry.

### Effect of infection rate on vomocytosis

Dendritic cells and MΦs were seeded on 24 well plates (75,000 cells/ well [DCs] and 50,000 cells/ well [MΦs]) in complete media and incubated at 37°C and 5% CO_2_. Infections with CNs and subsequent time-lapse imaging studies were performed between day 11-15. Wildtype CN H99 and the acapsular mutant *cap59* CN (both generously gifted by Dr. Angie Gelli, UC Davis, CA) were grown first on yeast extract peptone dextrose (YPD) agar (Thermo Fisher Scientific, Waltham, MA) followed by transferring a single colony to YPD broth (Thermo Fisher Scientific) shaking at 30°C overnight. The next day, CNs were washed with PBS (x3) via centrifugation. The heat-killed (HK) CN negative control group was prepared by incubating CNs at 70°C on a heat block for 1 hour. Both live and HK CNs were opsonized with 10 ug/ml of the anti-capsular IgG1 monoclonal antibody 18B7 (supplied from both Sigma and the Casadevall Lab, Johns Hopkins University, MD) and 50% human AB serum (Sigma). Opsonized pathogen was co-incubated with phagocytes at a 1:1 or 5:1 CN:phagocyte ratio (c.p.r.) for 2 hours in media with 10% human AB serum. Next, infected culture wells were washed with complete media (x5) to ensure all extracellular CNs were removed. Lastly, complete media was added, and time-lapse imaging was performed on infected cells for 14 hours.

### Effect of Drugs on Vomocytosis

For drug-treated experimental groups, DCs were infected with CN at a 5:1 c.p.r. using identical methods outlined previously. After the 2-hour phagocytosis and washing step, the wells were treated with DC media containing either 10uM of chloroquine [CQ] (Thermo Fisher Scientific), 100 nM of cytochalasin B from *Drechslera dematioidea* [CYT low] (Sigma), 4uM of cytochalasin B (CYT hi), or 100 nM of bafilomycin A1 [BFA] (Sigma). These parameters were selected based on the range of drug concentrations used in prior vomocytosis studies^15,17,24,44^. This step was followed by time-lapse imaging for 14 hours whilst in drug-containing media.

### Effect of Polarizing Agents on Vomocytosis

For experiments studying the effect of phagocyte polarization on vomocytic frequency, DEX or LPS were added to DCs at 48 hours prior to CN infection. For MΦs, IL4/13 or LPS was added to the cells 48 hours prior CN infection. Polarizing agent-containing media was replaced with fresh complete media prior to infection with CN. The infection then proceeded using identical methods as previously outlined.

### Time-lapse Imaging

Infected cells were kept at 37°C and 5% CO_2_ in the imaging chamber of the BZ-X Fluorescence Microscope (Keyence, Itasca, IL). Images were taken every 4 minutes for a period of 14 hours and compiled into a single movie file using BZ-X software. Movies were blinded by a third party before manual tracking of CNs and scoring for vomocytosis events by an independent technician.

### Confocal Time-Lapse and High-Resolution Imaging

Prior to the experiment, DCs were stained with 1,1’-Dioctadecyl-3,3,3’,3’-Tetramethylindodicarbocyanine, 4-Chlorobenzenesulfonate Salt (DiD; 1µM; Thermo Fisher Scientific) and CN cells were stained with FITC-NHS (1mg/ml). Three hours after infection, cells were placed in the imaging chamber of Olympus FV3000 (Olympus Corporation, Westborough, MA) at 37°C and 5% CO_2_. Images were taken every 10 minutes for the period of 5 hours at 20x magnification and analyzed for vomocytosis events.

For confocal high-resolution imaging, DCs were stained with DiD (1µM) and CN were stained with Calcofluor White (CFW; 1mg/ml; Sigma). After 2 hours of infection at a 1:1 c.p.r., the cells were washed 5 times with DC media. Next, 4 hours after washing, the sample was fixed with 2% paraformaldehyde and vomocytosing cells were imaged using the Olympus FV3000 confocal at 60x magnification.

### DC viability

Dendritic cells were seeded on 24-well plates (75,000 cells/ well) in DC media and incubated at 37°C and 5% CO_2_. Cells were co-incubated with CNs opsonized using previously mentioned methods. Extracellular CNs were washed with DC media and 14 hours later cell viability was measured using the CyQUANT lactate dehydrogenase (LDH) cytotoxicity assay (Thermo Fisher Scientific) according to manufacturer’s instructions. Additional details on the viability assays are provided in the Supplementary Information.

### Data and Statistical Analysis

In each experimental group replicate, 300 randomly selected CNs from multiple viewing regions were observed and vomocytosis manually quantified. All statistical analyses were performed using GraphPad Prism 9. Categorical data of vomocytosis frequency in the different conditions was assessed by one-way ANOVA corrected for multiple comparisons by false discovery rate (FDR) using a two-stage linear step-up procedure of Benjamini, Krieger and Yekutieli (GraphPad® Prism). This same statistical analysis procedure was used to evaluate flow cytometry marker expression data, number of CN cells per phagosome data, and timing of expulsion data. For F4/80 and CD11c expression comparison, an unpaired t-test was used as there were only two groups to compare. All data shown include at least three independent experiments. Original time-lapse movies, upon which manual scoring was performed, are freely available upon request. All column graphs, generated on GraphPad Prism 9, display the individual data points, mean, and standard error mean (SEM). All violin plots, generated on R, visualize the individual event data points, mean (red dot), and box plot containing median and interquartile range.

## Results

### DC and MΦ phenotype validation

The phenotypes of the bone-marrow derived DCs and MΦs were characterized via flow cytometry to confirm that the cell cultures generated with these protocols were authentic. These cells were characterized for the DC marker - CD11c, and the MΦ marker - F4/80, via flow cytometry. (**Figure 1A**) The purity of MΦ and DC cultures based on presence or lack of F4/80 and CD11c markers are visualized via bar graphs. Dendritic cell cultures exhibit low F4/80+ and high CD11c+ gated populations, while MΦ cultures display high F4/80+ and low CD11c+ gated populations. (**Figure 1B**) These results confirm that the cultures grown are genuine bone marrow-derived DCs and MΦs^45,46^.

**Figure 1.**
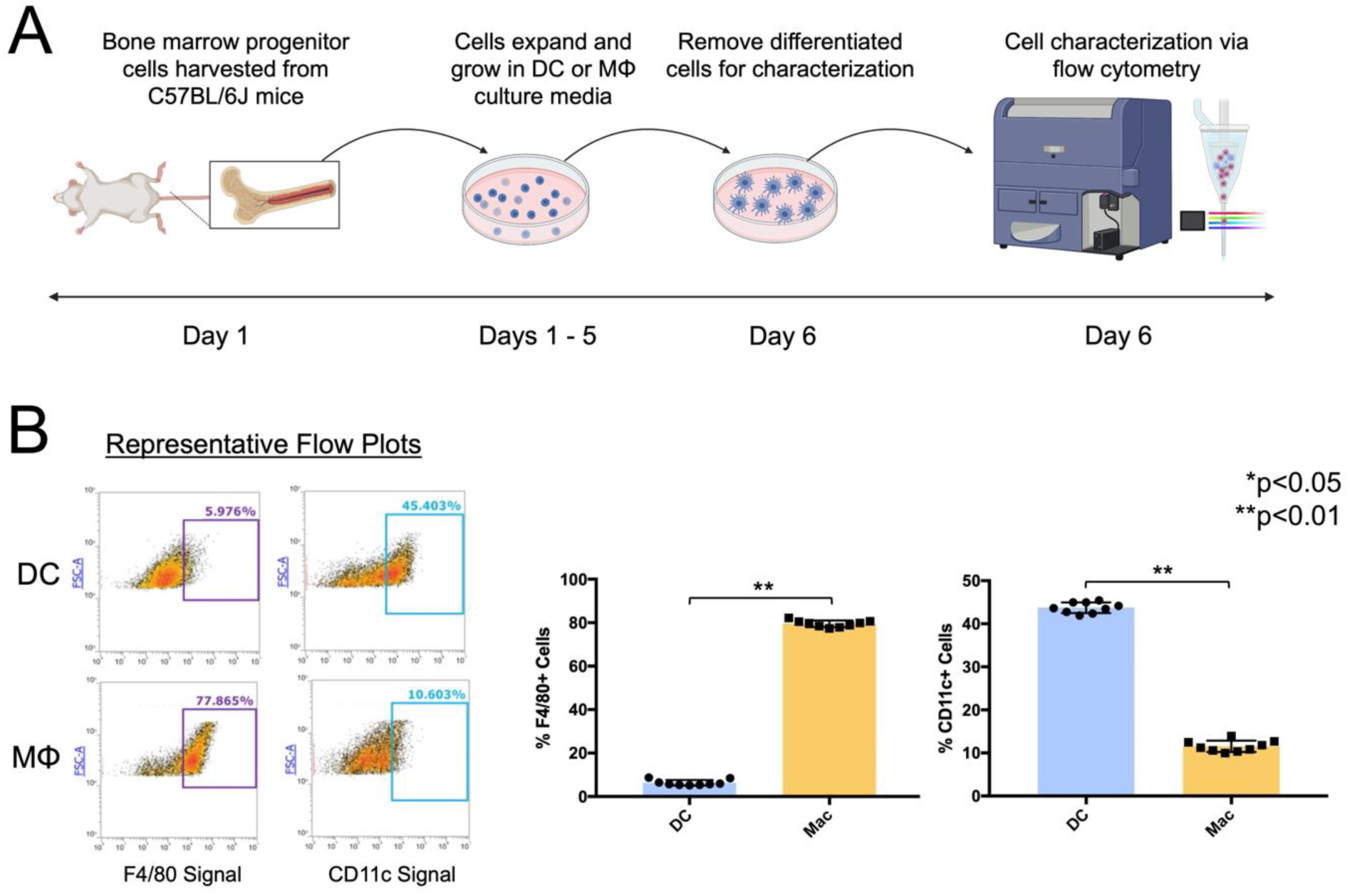
Verification of DC and MΦ phenotypes via F4/80 (MΦ marker) and CD11c (DC marker) staining and flow cytometric analysis. (**A**) Schematic of flow cytometry experiment. Bone marrow progenitor cells were obtained from C57BL/6J mice and grown in either DC differentiation media (GM-CSF supplemented) or MΦ differentiation media (M-CSF supplemented via L929). On day 6, these cells were stained and analyzed via flow cytometry. (**B**) Flow cytometry data for F4/80 and CD11c identification markers. The flow populations were gated to determine the percentage of cells exhibiting high expression of each marker. Representative flow cytometry plots of differentiated DCs and MΦs are shown. Column graphs of full flow cytometry data show the percentage of cells positive for F4/80 and CD11c for DC-differentiated and MΦ-differentiated cultures (N=3, n=9, statistical analysis performed using an unpaired t-test).

### Vomocytosis from DCs is independent of CN capsule and infection rates, and is comparable to vomocytosis from MΦs

We quantified vomocytic events using time-lapse microscopy and verified their non-lytic nature via cell viability assays. After 2-hour phagocytosis of CN, phagocytic cells were washed for the removal of extracellular CN, and time-lapse imaging experiments were performed during a period of 14 hours (**Figure 2A**). Vomocytosis events, defined as expulsions of CN from host cell while both remain intact, were observed at both CN:phagocyte ratios (c.p.r.) of 1:1 and 5:1, as shown in representative time lapse images (**Figure 2B-C**). Confocal time-lapse microscopy videos confirmed vomocytosis events by visually verifying instances of increased CN (green) fluorescent intensity upon uncoupling between DC and CN cells over the course of 8 hours (**Figure S1**). Overall, vomocytosis occurred at a rate of 17% for MΦs infected at a 1:1 c.p.r., 18% for MΦs infected at a 5:1 c.p.r., 11% for DCs infected at a 1:1 c.p.r., 13% for DCs infected at a 5:1 c.p.r., and 13% for DCs infected with *cap59* CN at a 5:1 c.p.r. (**Figure 2D-E**). For the HK CN control groups, an expulsion rate of only 2% or less was observed for both DC and MΦ groups infected at 1:1 and 5:1 c.p.r. To confirm that these events were indeed non-lytic, DC viability was tested via an LDH assay. We observed no increased toxicity due to CN infection (**Figure S2**). Notably, the vomocytosis rates observed from DCs were insignificant to those observed from MΦs at 1:1 and 5:1 c.p.r. (**Figure 2F**). For conclusive visual confirmation of vomocytosis, high resolution confocal microscopy was used to observe two fixed DC cells mid-vomocytosis at 6 hours after infection with CN (**Figure 2G**).

**Figure 2.**
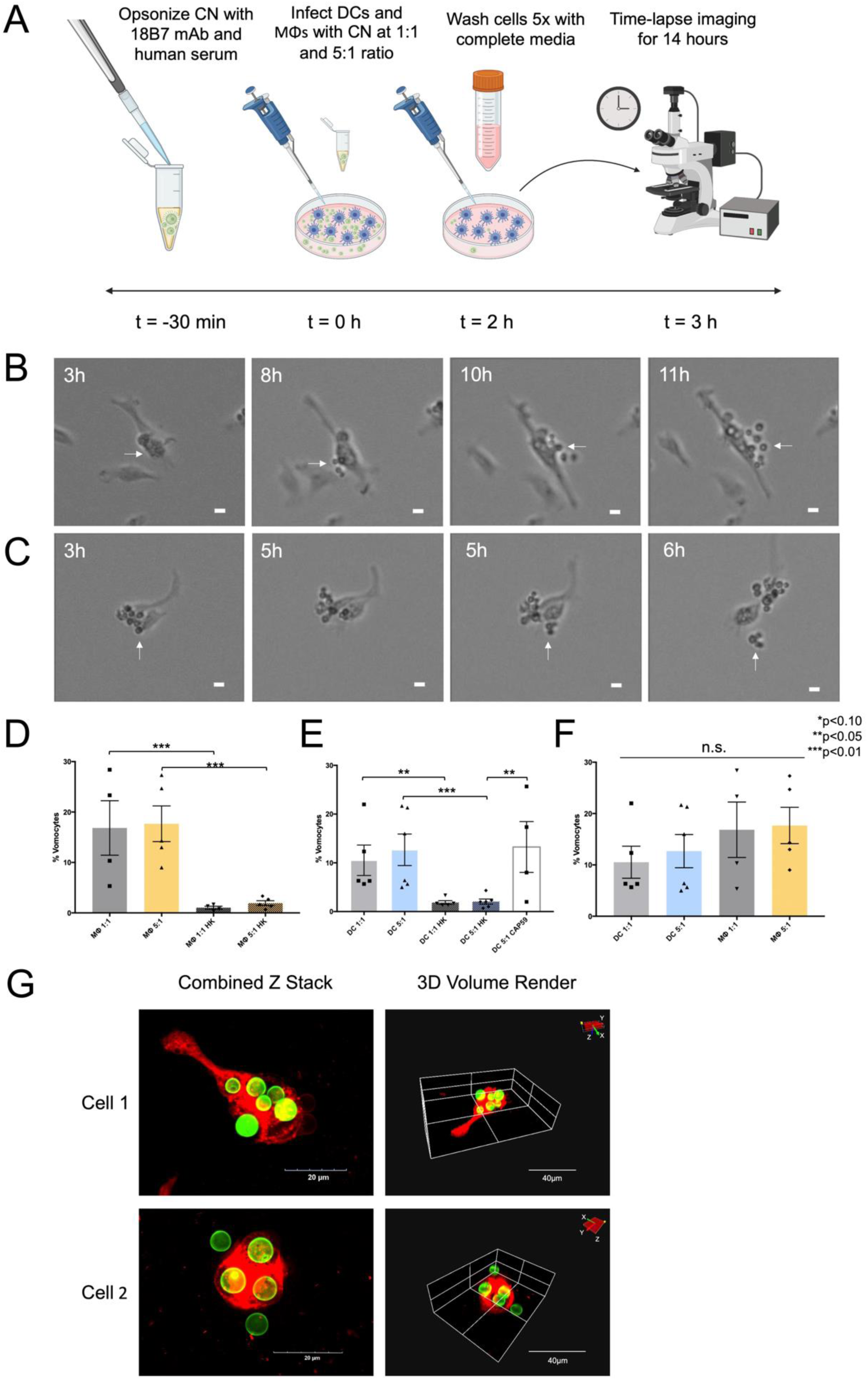
Time-lapse analysis of vomocytosis from DCs. (**A**) Schematic of time-lapse experiment. CN was prepared for phagocytosis by opsonizing with 18B7 mAb and human serum. DCs or MΦs were infected with opsonized CN either at a 1:1 or 5:1 c.p.r. for 2 hours. Following infection, the phagocytes were washed 5x and time-lapse imaged for 14 hours. Representative time-lapse images of DCs performing vomocytosis at (**B**) 1:1, or (**C**) 5:1 c.p.r. (scale bar = 10µm). (**D-F**) Graphs of vomocytosis rates of MΦs and DCs are shown at 1:1 and 5:1 c.p.r., compared to a HK CN control and acapsular *cap59* CN (N≥4 for each condition, statistical analysis performed using one-way ANOVA corrected for multiple comparisons by FDR using a two-stage linear step-up procedure of Benjamini, Krieger and Yekutieli). **(G)** Confocal images showing instances of DCs (DiD, red) expelling CN (CFW, green). The events are visualized via both a combined Z stack and a 3D volume rendering.

### Drugs that disrupt phagolysosomal maturation and actin polymerization inhibit vomocytosis

Next, DCs were treated with drugs previously documented to affect phagolysosomal maturation, and vomocytosis from MΦs. Namely, CQ, CYT, and BFA were used for these experiments (**Figure 3A**). After assessing potential toxicity to DCs and CNs at the desired concentrations (**Figure S2**), the individual drugs were added to wells containing CN-infected DCs at a 5:1 c.p.r., just prior time-lapse imaging. All drugs (at the selected concentrations) reduced vomocytosis rates (**Figure 3B**). For DCs, we observed vomocytosis rates of 4%, 5%, 6%, and 6% for CQ, CYT hi, CYT lo, and BFA respectively. Vomocytosis rates in both, CQ- and CYT hi-treated cells were not significantly different from the rate observed for the HK CN group. We confirmed that these inhibited rates were not due to the impact of the drugs on immune cell and CN cell viability (**Figure S3**). Furthermore, in CN-infected DC cultures treated with these drugs, there were no observed increases in toxicity to extracellular or intracellular CNs (**Figure S4**).

**Figure 3.**
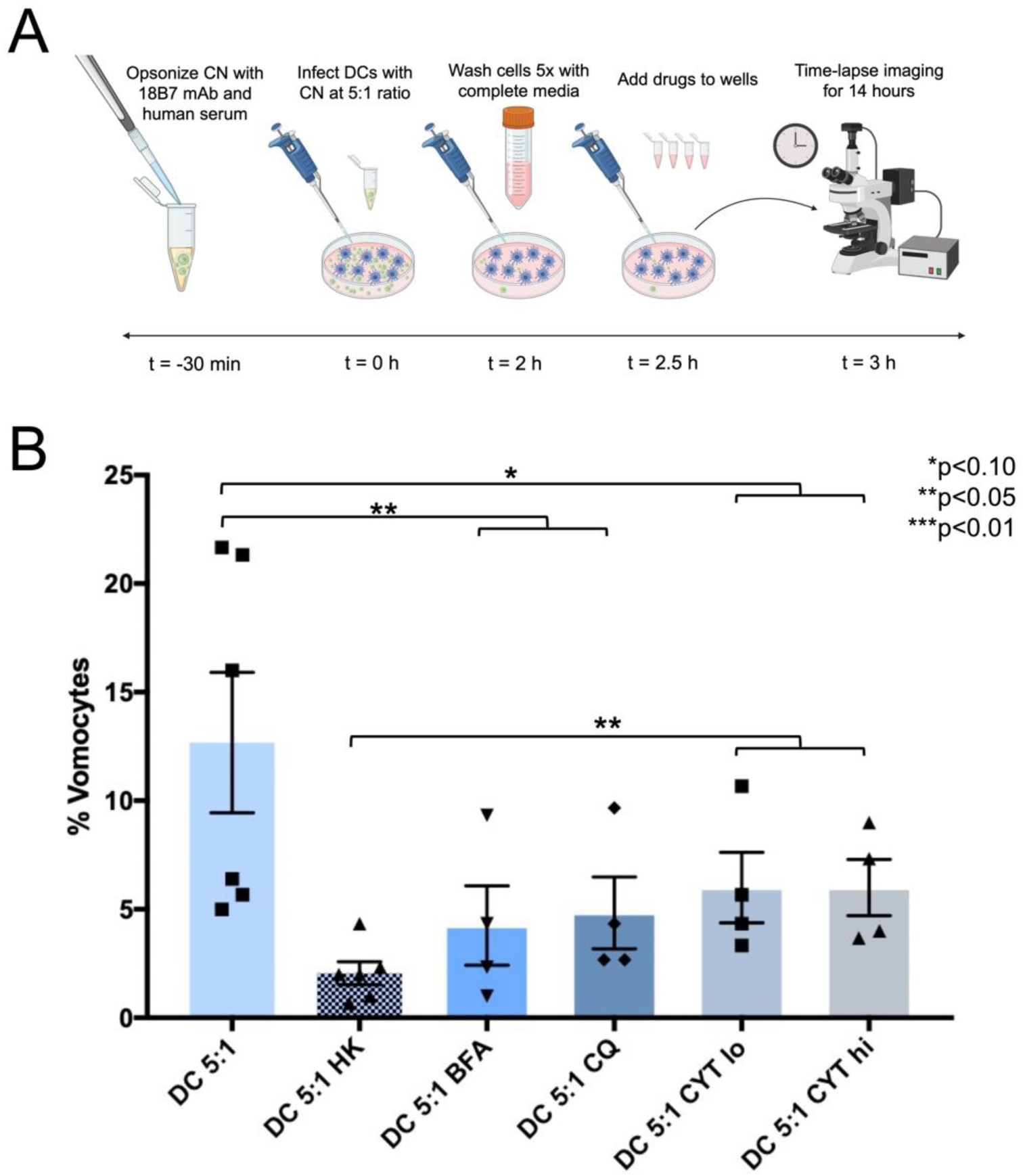
Time-lapse analysis of DC vomocytosis rates under exposure to vomocytosis-modulating drugs. (**A**) Schematic of time-lapse experiment. CN cells were prepared for phagocytosis by opsonizing with 18B7 mAb and human serum. DCs were infected with opsonized CN at a 5:1 c.p.r. for 2 hours. Following infection, the wells were washed 5x, replaced with drug-containing media, and time-lapse imaged for 14 hours. (**B**) Vomocytosis rates of DCs, at a 5:1 infection ratio, are shown in untreated, HK CN, and drug-treated conditions—either BFA, CQ, CYT lo, or CYT hi (N≥4 for each condition, statistical analysis performed using one-way ANOVA corrected for multiple comparisons by FDR using a two-stage linear step-up procedure of Benjamini, Krieger and Yekutieli).

### Treatment of MΦs and DCs with polarization agents alters their immune phenotype

Prior to testing the effect of immune state on vomocytosis, verification of the immunophenotype of MΦs and DCs was performed (**Figure 4A**). In these experiments, the immature MΦ (iMΦ; Control), activated (LPS), anti-inflammatory (IL4/13), and tolerized then challenged (IL4/13 + LPS) conditions were probed (**Figure 4B**). The LPS-treated group displayed significantly higher inflammatory marker expression of CD38 and iNOS compared to the immature, untreated group. Interestingly, the tolerogenic marker Arg1 was also increased on the LPS-treated MΦs. On the other hand, the IL4/13-treated group showed no significant differences for CD38 and iNOs, compared to the immature group. Whilst exhibiting higher Arg1 expression than the untreated and LPS groups. Additionally, the tolerogenic group challenged with LPS displayed similarly high levels of Arg1. Additionally, this group had similar expression of CD38 and iNOS markers to the LPS-treated cells.

**Figure 4.**
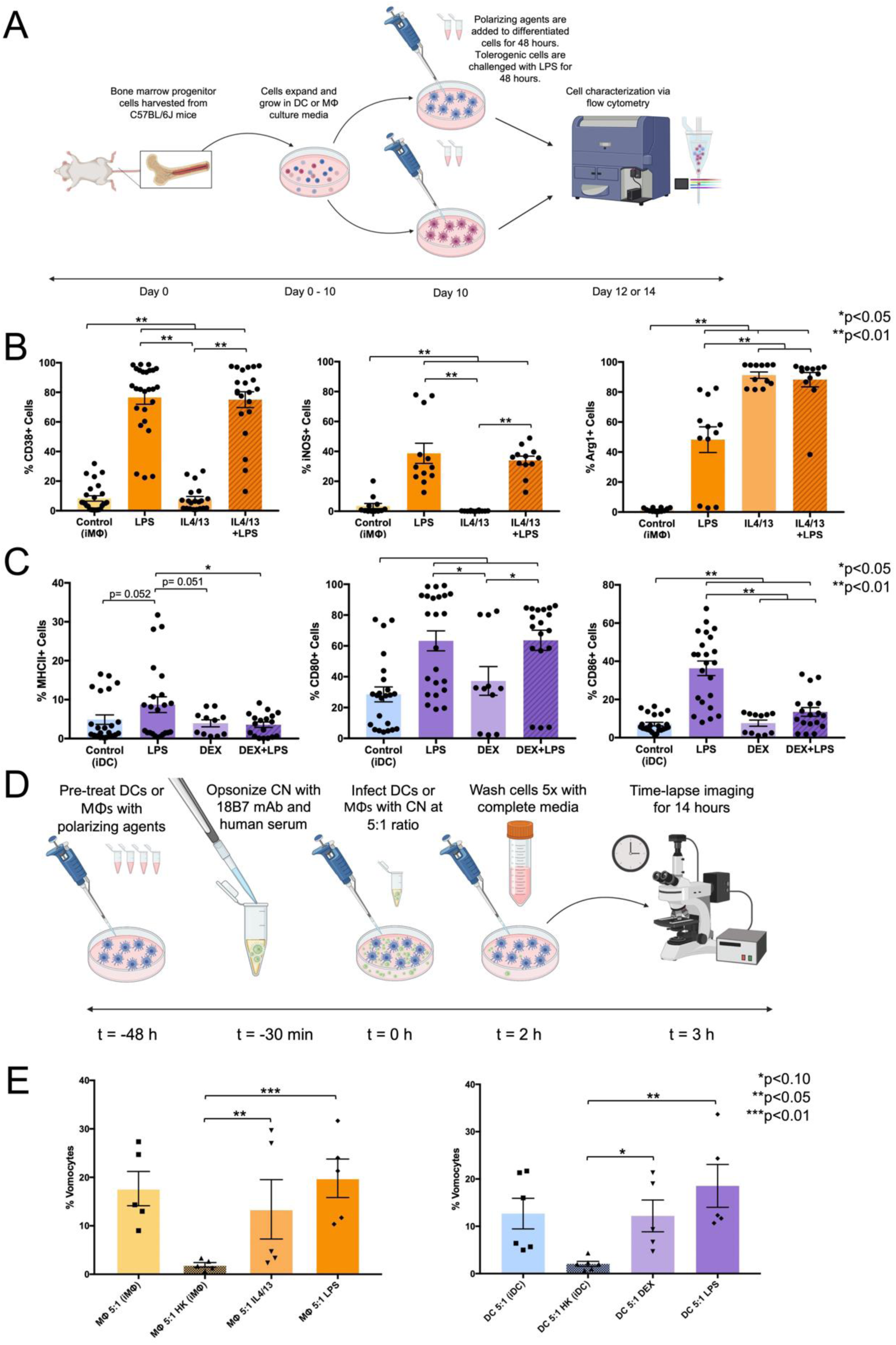
DC and MΦ vomocytosis rates following immune polarization. (**A**) Schematic of flow cytometry experiment. Dendritic cells and MΦs were incubated with polarization agents for 48 hours. Dendritic cells were polarized using dexamethasone, or LPS for tolerized or inflammatory phenotypes, respectively. Macrophages were polarized using agents LPS or IL4/13 for inflammatory (M1) or tolerogenic (M2) phenotypes, respectively. Also, an additional experimental group of tolerized DC and MΦ were challenged with LPS for another 48 hours. These groups were stained for DC immunophenotype markers (MHCII, CD80, and CD86) or MΦ immunophenotype markers (CD38, iNOS, and Arg1) and analyzed via flow cytometry. (**B**) Confirmation of DC immunophenotype characterization by flow cytometry readout of maturation markers MHCII, CD80, and CD86. (**C**) Confirmation of MΦ immune phenotype characterization by flow cytometry readout of inflammatory activation markers CD38 and iNOS, as well as the M2 marker Arg1. (**D**) Schematic of time-lapse experimental design. Briefly, prior to infection, DCs and MΦs were incubated with polarization agents for 48 hours. Then, CN was prepared for phagocytosis by opsonizing with 18B7 mAb and human serum. Polarized DCs and MΦ were infected with opsonized CN at a 5:1 c.p.r. for 2 hours. Following infection, the phagocytes were washed 5 times and time-lapse imaged for 14 hours. (**E**) Vomocytosis rates of DCs and MΦs at a 5:1 infection ratio are shown for immature (untreated), HK CN, tolerized, and inflammatory states. (N≥4, n≥11 for each condition, statistical analysis performed using one-way ANOVA corrected for multiple comparisons by FDR using a two-stage linear step-up procedure of Benjamini, Krieger and Yekutieli).

For DCs, the inflammatory markers MHCII, CD80, and CD86 were analyzed via flow cytometry on CD11c+ gated cells. The immature DC (iDC; Control), activated (LPS), and tolerized (DEX) conditions were analyzed. Additionally, an added DEX-treated DC group was challenged with LPS (DEX + LPS) for an additional 48 hours to test resistance to inflammatory activation. The LPS-treated group displayed significantly higher MHCII, CD80, and CD86 expression compared to the immature untreated group (**Figure 4C**). Meanwhile, the DEX group displayed no significant difference to the immature group on the basis of these inflammatory markers. Furthermore, when tested with LPS after DEX treatment, these tolerogenic DCs displayed significantly lower activation of MHCII and CD86 expressions compared to the LPS-treated group, indicating resistance toward maturation.

### Immune polarization of MΦs and DCs does not affect vomocytosis rate

Next, the vomocytosis rates of polarized MΦs and DCs were tested (**Figure 4D**). For MΦs, the anti-inflammatory (IL4/13) and pro-inflammatory (LPS) conditions showed significantly higher vomocytosis rates than the heat killed control. However, there were no significant differences in rates between the untreated (18%), anti-inflammatory (13%), or pro-inflammatory (20%) MΦs(**Figure 4E, left**). Similarly, DCs polarized to tolerogenic (DEX) and inflammatory (LPS) phenotypes displayed a higher vomocytosis rate than that of the HK CN group. However, the DC vomocytosis rates of the tolerogenic (12%) and inflammatory (19%) groups were not significantly different to each other or the untreated group (13%) (**Figure 4E, right**).

### Infection Ratio, Drug treatments, and Immune Polarization affect Vomocytosis Kinetics

We used time-lapse videos to measure of exact time of expulsion for each vomocytic event. There was no difference in vomocytosis timing for DCs infected with CN at a 1:1 and 5:1 c.p.r., and no difference compared to DCs infected with the *cap59* CN (**Figure 5A**). For MΦs, the 5:1 c.p.r. condition displayed significantly lower average time to expulsion compared to the 1:1 c.p.r. group (**Figure 5B**). Comparing DCs and MΦs, the MΦ 1:1 c.p.r. condition showed a higher time to expulsion than both the DC 1:1 and DC 5:1 c.p.r. conditions (**Figure 5C**). For the drug-treated DC groups, both the CQ and CYT hi treatments performed vomocytosis faster than the DC 5:1 untreated control (**Figure 5D**). Further, both immune-polarized DCs showed significantly lower time of expulsion than the unpolarized control, with the LPS-treated group having shorter expulsion times than the DEX-treated group (**Figure 5E**). In MΦs, both polarized conditions displayed insignificant differences in timing compared to the control. However, the LPS-treated condition had a significantly faster time of expulsion than the IL4/13-treated group (**Figure 5F)**.

**Figure 5:**
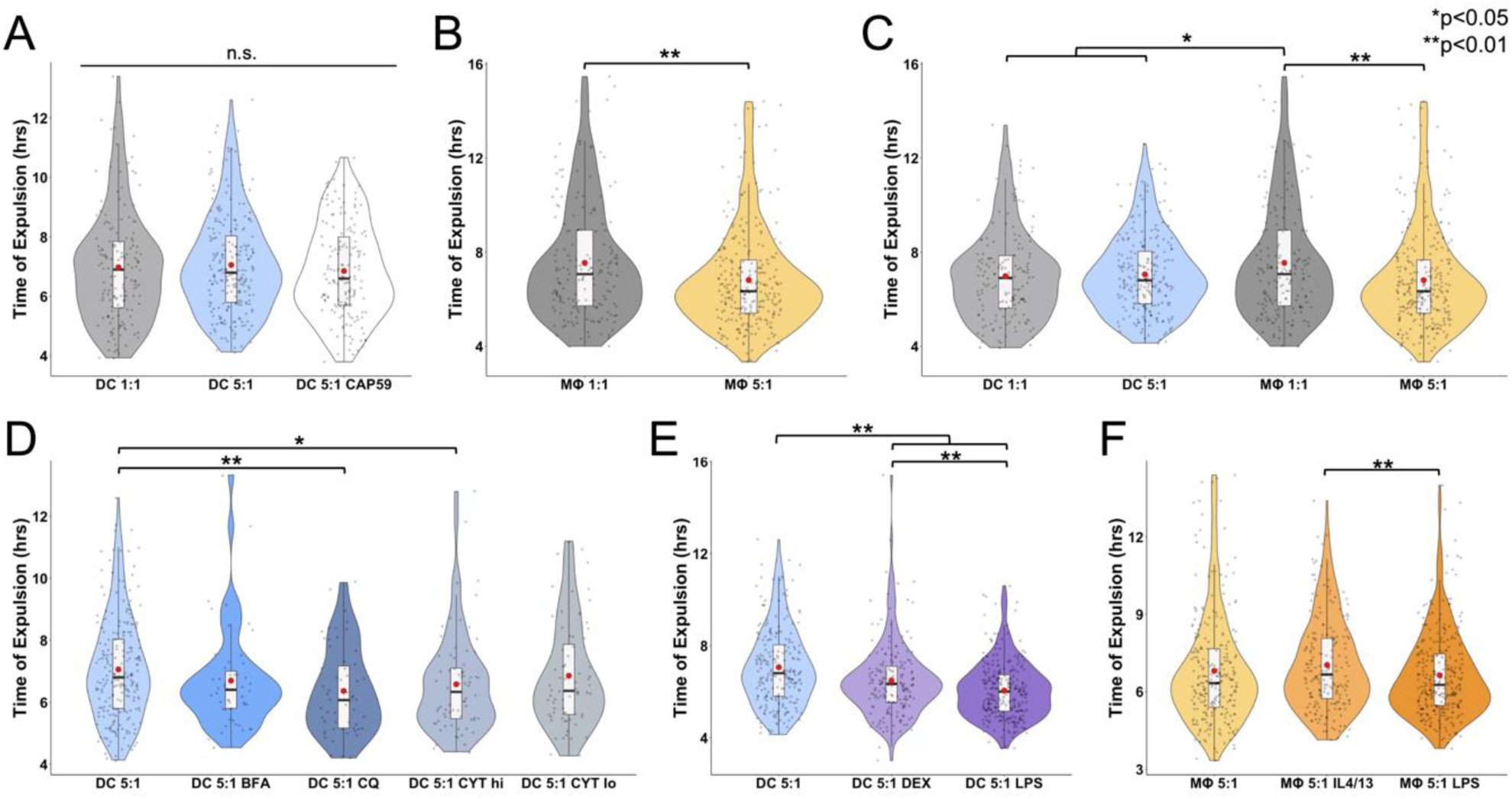
Vomocytosis event timing analysis. Results are displayed in violin box plots with individual dots representing each timing event. The red circle in each plot represents the mean timing occurrence. (**A**) Violin plot displaying timing of DC vomocytosis events compared between 1:1 and 5:1 c.p.r., as well as 5:1 c.p.r. with *cap59* CN. (**B**) Violin plot of MΦ vomocytosis timing between 1:1 and 5:1 c.p.r. (**C**) violin plot comparing vomocytosis timing between DC and MΦ at different CN infection ratios. D) Violin plot of DC vomocytosis expulsion timing under a 5:1 infection ratio of CN under different drug treated conditions (BFA, CQ, CYT lo, or CYT hi) compared to an untreated control. (**E**) Violin plot of DC vomocytosis timing under a 5:1 CN infection as an immature phenotype (untreated), tolerogenic phenotype (DEX treated), and inflammatory phenotype (LPS treated). (**F**) Graph of MΦ vomocytosis timing under a 5:1 CN infection as an immature M0 phenotype (untreated), tolerogenic M2 phenotype (IL4/13), and inflammatory M1 phenotype (LPS treated) (N≥4, n≥51 for each condition, statistical analyses were performed using one-way ANOVA corrected for multiple comparisons by FDR using a two-stage linear step-up procedure of Benjamini, Krieger and Yekutieli).

### Different treatments affect number of CN per vomosome

In addition to characterizing the vomocytosis rates and timing of different conditions, the number of CN located in the phagosome before a vomocytosis event, or ‘vomosome’, was also documented and analyzed. Under different infection ratios, the 5:1 condition had a significantly higher average number of CN per vomosome compared to the 1:1 infection ratio for DCs. Additionally, the 5:1 c.p.r. *cap59* CN mutant condition displayed a higher number of CN per vomosome than both the 5:1 and DC 1:1 c.p.r. for DCs. (**Figure 6A**) However, in MΦs there was no significant difference in CN per vomosome between the 1:1 and 5:1 infection ratios (**Figure 6B**). Notably, both the 1:1 and 5:1 c.p.r. MΦ groups showed higher number of CN located in their vomosomes than both DC 1:1 and DC 5:1 c.p.r. groups (**Figure 6C**). When analyzing drug-treated conditions, the DC 5:1 c.p.r. groups treated with CQ, CYT hi, or CYT lo had a significantly higher number of CN per vomosome than the control. The BFA-treated DC group showed no significant difference in CN per vomosome compared to the untreated control (**Figure 6D**). Finally, immune polarized groups were analyzed for differences in number of CN per vomosome. Both DEX-treated and LPS-treated DCs were observed to have a higher number of CN located in the phagosome prior to a vomocytosis event compared to the unpolarized control. Additionally, the LPS-treated DC group had a higher CN per vomosome count than the DEX-treated group (**Figure 6E**). For polarized MΦs, IL4/13-treated cells displayed a lower CN per vomosome count to the control. Conversely, the LPS-treated group had a higher CN per vomosome average than the control, as well as IL4/13-treated groups. (**Figure 6F**).

**Figure 6:**
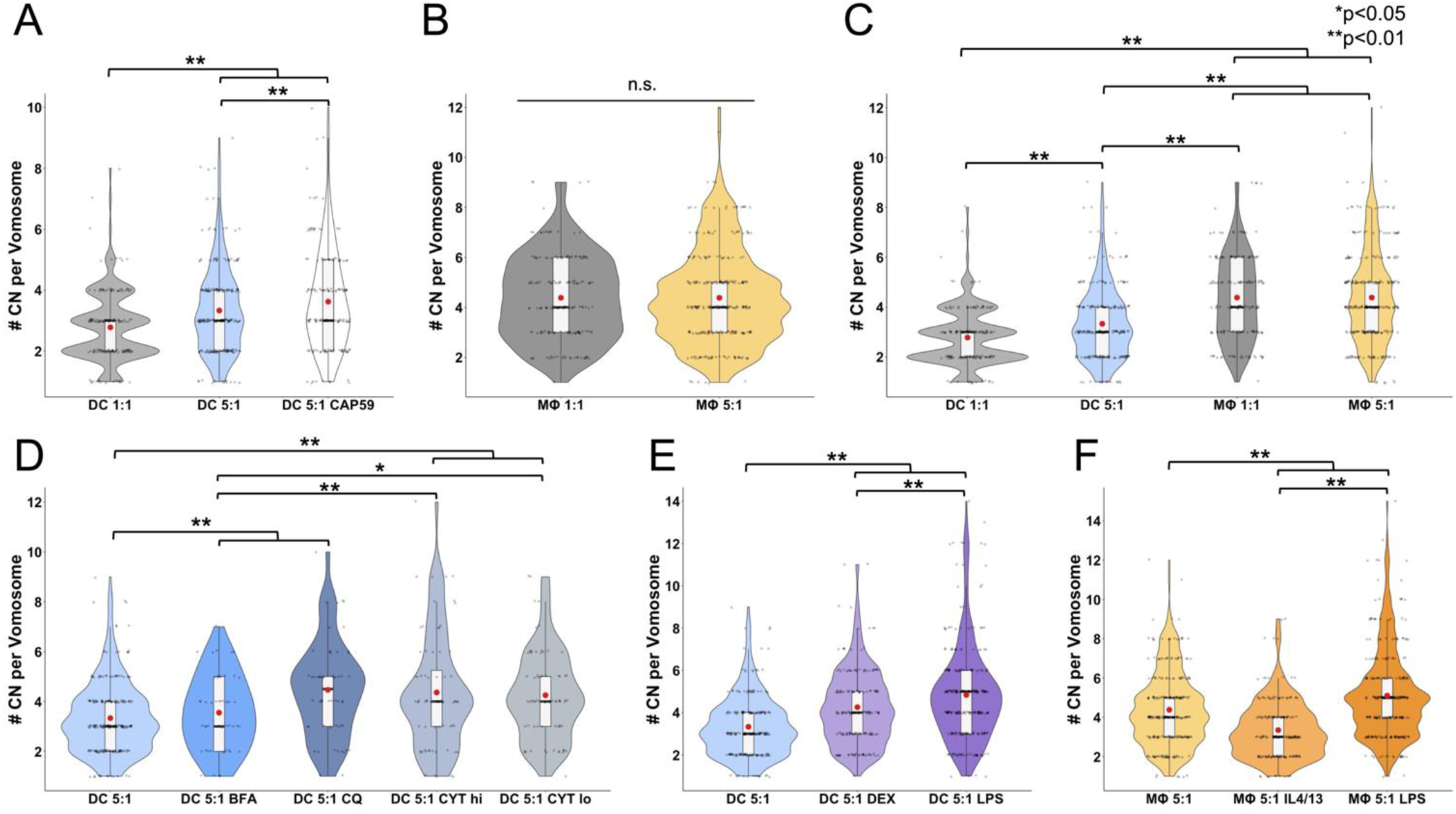
Number of CN per vomosome. Results are displayed in violin box plots with individual dots representing # CN per vomosome for each event. The red circle in each plot represents the mean # CN for each condition. (**A**) Violin plot displaying # CN per DC vomosome compared between 1:1 and 5:1 CN infection ratios, as well as 5:1 infection with a *cap59* CN. (**B**) Violin plot of # CN per MΦ vomosome compared between 1:1 and 5:1 CN infection ratios. (**C**) Violin plot comparing # CN per vomosome between DC and MΦ at different CN infection ratios. (**D**) Violin plot of # CN per vomosome for DCs with a 5:1 CN infection ratio under different drug treated conditions (BFA, CQ, CYT lo, or CYT hi) compared to an untreated control. (**E**) Violin plot of # CN per vomosome for DCs with a 5:1 CN infection as an immature phenotype (untreated), tolerogenic phenotype (DEX treated), and inflammatory phenotype (LPS treated). (**F**) Graph of # CN per vomosome for MΦs with a 5:1 CN infection as an immature M0 phenotype (untreated), tolerogenic M2 phenotype (IL4/13-treated), and inflammatory M1 phenotype (LPS treated) (N≥4, n≥51 for each condition, statistical analyses were performed using one-way ANOVA corrected for multiple comparisons by FDR using a two-stage linear step-up procedure of Benjamini, Krieger and Yekutieli).

## Discussion

Our novel results show that live CN can perform vomocytosis from DCs, to a similar extent exhibited by both MΦs and neutrophils. Here, the occurrence of this phenomenon from DCs was rigorously verified by time-lapse microscopy, confocal imaging, and viability assays. Additionally, the average rate and timing of vomocytosis events from DCs were the same as those from MΦs. Given our results on DC vomocytosis and those published on MΦs and neutrophils, it is likely that some components of this mechanism may be conserved across these cell types derived from common myeloid progenitor cells.

Vomocytosis rates from DCs were observed to have no significant correlation to infection rate or presence of CN capsule. However, DCs infected with *cap59* CN contained a significantly higher number of CN per vomosome compared to DCs infected with the wildtype strain (H99). This is not surprising, as the CN capsule is known to be a potent anti-phagocytic agent that plays a role in reducing uptake by MΦs^47–49^ and DCs^50^. Additionally, DCs infected at a 5:1 ratio showed higher amount of CN per vomosome than DCs 1:1 infected, which may be attributed to the larger number of CN that the DCs encountered during the experiment.

When treated with drugs to modify the phagosome (CQ, BFA) or actin polymerization (CYT lo, CYT hi), DCs displayed substantially lower rates of vomocytosis compared to the untreated control. Chloroquine is a weak base that passively diffuses into acidic organelles in the cytoplasm, becomes protonated and prevents maturation and fusion of endosomes and lysosomes^51,52^. Cytochalasin B is an actin polymerization inhibitor that has been demonstrated to induce a release of lysosomal enzymes, modulating lysosomal fusion with phagosomes^53,54^. Lastly, BFA is involved in the inhibition of the vacuolar ATPase in lysosomes^55^. With respect to CQ, our findings conflict with previous studies that have successfully demonstrated this drug to increase vomocytosis from J774 MΦ cell lines^10,17^. However, *Yang et al* showed that CQ decreases vomocytosis rates from primary murine neutrophils^24^. These discrepancies could be attributed to differences in cell type, as well as source (cell line vs. primary murine cells). Interestingly, the decreased BFA-treated vomocytosis rates we observed from DCs align with observations by *Nicola et al* (from J774 macrophage cells), that this drug treatment reduces vomocytosis rates^17^. On the other hand, neutrophils were shown to have no change in vomocytosis rates after BFA treatment^24^. Again, these differences could be due to cell type, as well as cell source. Overall, our findings indicate that vomocytosis from DCs is indeed inhibited by phagosomal alkalinization— by either the weak base CQ or the ATPase inhibitor BFA. Pertaining to CYT treatment, again there is some disagreement within literature on the effect of this drug on vomocytosis. *Dragotakes et al* observed a decrease in vomocytosis rates of J774 MΦs following treatment with either cytochalasin B or D^44^. Conversely, *Alvarez et al*. saw increased rates of expulsion in the same cell type following cytochalasin D treatment^9^. In neutrophils, cytochalasin D was also seen to increase vomocytosis rates in a study by *Yang et al*^24^. Our observations for DCs align with those published by *Dragotakes et al*., but are in conflict with the other two studies. It is possible that cytochalasin B may cause lysosomal fusion in the infected DCs and cripple the CN’s ability to thrive in the phagosome, although our viability data suggests that intracellular CN are still intact during this treatment. The role of CYT in inhibiting actin polymerization may play a more significant role in limiting the phagocyte’s machinery and preventing CN from inducing an expulsion. In investigating average timing of vomocytosis, the CQ-treated and CYT hi-treated vomocytosis events displayed a statistically shorter length of time before expulsion compared to the untreated group. Chloroquine and CYT both reduce expulsion rates, but the CN are released relatively quicker compared to untreated cells. These events may be due to alternate mechanisms such as endosome recycling pathways^56^. Interestingly, within the drug treated groups, the average number of CNs per vomosome were all significantly higher than the untreated group with the exception of the BFA-treated group. It is possible that BFA alkalinization of the phagosome reduces CN dividing ability, as CN has been observed to replicate faster in acidic environments^18,57^.

Treatment of MΦs and DCs with polarization agents did not affect the vomocytosis rates compared to the unpolarized control. This result is contrary to some documented literature for MΦs, as *Gilbert et al*. found that inflammatory-polarized MΦs (via ERK5 inhibition) displayed a higher rate of expulsion than untreated MΦs, shown in both primary human MΦs and J774 cells^21^. Other conflicts exist where *Voelz et al*. showed that anti-inflammatory IL-4 or IL-13 treated MΦs perform a lower rate of vomocytosis compared to an untreated control, also in both primary human MΦs and J774 cells^23^. However, the same study showed that the pro-inflammatory IFN-γ, TNF-α, or IL-17 treated human and J774 MΦs did not have a significantly different vomocytosis rate than unpolarized MΦs, which aligns with our observations. Interestingly, a recent study by *Zhang et al*. found that primary murine MΦs treated with inflammatory extracellular vesicles displayed lower vomocytosis rates^58^— this result contradicts both this study and previous studies. Overall, investigation of the effect of immune phenotype on vomocytosis has been inconclusive. This may be attributed to differences in cell types, polarization procedures, phenotype validation and vomocytosis observation methods. Our results suggest that immune polarization is not a significant influence on vomocytosis rates from MΦs or DCs. However, there was a significant difference in the average time for vomocytosis occurrence for these groups. Cryptococcal cells performed vomocytosis in a significantly shorter time from LPS-treated DCs compared to DEX-treated DCs. Similarly, the average non-lytic exocytosis time of CNs in LPS-treated MΦs was significantly longer in time length compared to that of IL4/13-treated MΦs. This observation is likely due to the differences in molecular and physicochemical characteristics involved in phagolysosomal maturation between activated and tolerogenic phenotypes. For instance, M2 macrophages have been shown to undergo rapid and profound phagosomal acidification relative to M1 macrophages^59^. Moreover, the same study demonstrates that reactive oxygen species (ROS) production is much greater and more sustained in M1 than in M2 phagosomes. Perhaps the mature, ROS and cathepsin-rich phagosome of LPS-treated phagocytes creates an inhospitable environment for CNs, prompting them to escape at a faster rate than in the less harsh anti-inflammatory polarized phagosomes. Additionally, for both DCs and MΦs, the LPS-treated groups displayed higher numbers of CN per vomosome compared to the tolerogenic group. The higher number of CN per vomosome and faster ejection from LPS-treated DCs compared to DEX-treated DCs could suggest a direct correlation between number of CN and ejection time— this same trend was observed from DCs treated with CQ and CYT hi.

## Conclusions

In summary, this study documents the first recorded observation of DCs performing vomocytosis of CN. Moreover, multiple parameters were tested for their effects on vomocytosis including the infection ratio, presence of CN capsule, treatment with drugs, and polarization of phagocyte immune state. Vomocytosis from DCs is independent of infection ratio, CN capsule, or immune polarization phenotype. However, application of drugs that disrupt phagolysosome and actin processes significantly inhibit the DC vomocytosis. Interestingly, infection ratio, drug treatments, and immune polarization influence the timing of vomocytosis, as well as the number of CN present in the DC vomosome prior to expulsion. Overall, the capability of DCs to expel CNs following ingestion appears similar to that of MΦs based on occurrence rate, timing, and number of CN per vomosome. This finding could help to further elucidate infection dissemination mechanisms in immunocompromised patients, as this phenomenon is clearly conserved between multiple types of phagocytes, including DCs. Finally, more studies are needed to understand the mechanisms involved in vomocytosis from DCs, and moreover determine if there is commonality of processes across all the phagocytic cells that are implicated to perform this behavior. Further, CN is not the only pathogen that has been shown to induce vomocytosis. Therefore, questions on the cohesion of mechanisms across pathogens and phagocytes should be addressed in future investigations.

## Acknowledgements

We thank the Gelli Lab at UC Davis for providing the CN strains used in this study. Additionally, we thank the George Lab at UC Davis for graciously allowing us to use their confocal equipment and aiding in confocal imaging. Finally, we thank the Casadevall Lab for generously providing 18B7 antibody that was used for these experiments.

We acknowledge support from the following sources: NIH grant R35GM125012 (J.S.L.), NIGMS-funded Pharmacology Training Program grant T32GM099608 (N.P.), and NSF GRFP grant 1650042 (N.P.).

